# Microvascular pulsatility of the ageing brain and confounding effects of anaesthesia

**DOI:** 10.1101/2024.12.01.626169

**Authors:** Mia Viuf Skøtt, Elizaveta Melnikova, Eugenio Gutiérrez, Vladimir Matchkov, Leif Østergaard, Dmitry D Postnov

## Abstract

Microvascular pulsatility in the brain is crucial for sustaining the delicate balance between the brain’s metabolic demands and blood supply, as it affects nutrient exchange, waste removal and blood-brain barrier permeability. Abnormal pulsatility, associated with ageing or vascular risk factors, may impair the clearance of metabolic waste or contribute to conditions such as cerebral small vessel disease, ultimately leading to dementia and increasing the risk of stroke. In the present study, we introduce a new approach for robust full-field characterization of microvascular pulsatility and apply it to study cerebral perfusion and pulsatility across nearly the entire lifespan of awake and anaesthetized C57BL/6 mice. Our findings in awake animals reveal remarkably stable perfusion and pulsatility from 18 to 81 weeks, with pulsatility starting to change only in the final weeks of observation. In contrast, measurements taken under anaesthesia display a range of age-dependent and age-independent changes. We show that isoflurane affects the perfusion in an age-dependent manner, and both isoflurane and ketamine-xylazine, despite their distinct mechanisms, double the perfusion pulsatility and affect vascular diameter pulsatility in a complex manner, highlighting a previously overlooked detrimental effect of anaesthesia on assessing brain function.

## 1 Main

Microvascular pulsatility in the brain is crucial for sustaining the delicate balance between the brain’s metabolic demands and blood supply. The rhythmic oscillations in blood velocity and vessel diameters within the microvascular network contribute to efficient nutrient exchange and waste removal and affect the integrity of the blood-brain barrier. Normal cerebrovascular pulsatility has been identified as a key factor in facilitating fluid drainage from the interstitium to the perivascular spaces and thence to the extracerebral cerebrospinal fluid and venous system [1], while reduced pulsatility may impair the clearance of metabolic waste, contributing to the accumulation of toxic proteins such as amyloid-beta, a hallmark of Alzheimer’s disease [2]. Conversely, excessive pulsatility might cause mechanical stress on the vessel walls, leading to microvascular damage and contributing to conditions like cerebral small vessel disease, which, in turn, is strongly associated with cognitive decline and increased risk of stroke [3, 4]. While reduced cerebral blood flow has been widely proposed as the cause of small vessel disease, evidence suggests that vascular stiffening and associated changes in the pulsativity and autoregulation can be seen as an alternative key factor [5–9] as the persistent exposure of small vessels to strong pulsation may result in cerebral microvascular damage, leading to structural and functional changes in the brain. Ageing and vascular risk factors, such as hypertension and diabetes, are also associated with loss of arterial elasticity, which could lead to more pulsatile energy dissipating in the brain microvasculature and result in tissue damage [10, 11]. Furthermore, elevated pulsatility in the internal carotid or middle cerebral arteries has also been associated with white matter hyperintensities, indicators of cognitive decline and predictors of cognitive impairment and the development of dementia and stroke [12–14].

Characterising the transmission of cardiac pulsation through different brain regions and its impact on brain microvasculature is pivotal for understanding brain function and has broad clinical appeal. Several approaches and techniques have been used to quantify it in animal models and humans. Carotid-femoral pulse wave velocity is a gold standard measurement of arterial stiffness, but it only applies to central arteries and thus provides limited insight into downstream cerebrovascular beds [15]. Cerebral pulsatility indexes can be acquired with transcranial Doppler ultrasound [16–18] and time-of-flight or phase contrast magnetic resonance imaging [19], but they lack spatial resolution to access small vessels and are generally limited to large cerebral arteries, e.g. carotid or middle cerebral [20]. Recent advances in optical imaging have enabled more comprehensive imaging of microvascular pulsatility in animal models’ brain cortex [21] and human retina [22, 23]. Technologies like two-photon microscopy (TPM), optical coherence tomography, laser Doppler holography and laser speckle contrast imaging (LSCI) provide detailed, high-resolution images of microvascular dynamics in vivo [21–27]. These methods allow for the observation of pulsatility at both macroscopic and microscopic scales, offering deeper insights into the regional effects of pulsatility changes on brain tissue. Nevertheless, simultaneous full-field imaging of microvascular velocity and diameter pulsatility, crucial for understanding the complex function of the pulsating brain, remained a challenge.

In this study, we present a new algorithm and method for robust full-field characterisation of microvascular pulsatility in the brain cortex. We use laser speckle contrast imaging to acquire blood flow images at *≈* 200 frames per second, analyse them to achieve sub-frame sampling, and simultaneously extract pulsatility indexes for vessel diameters and blood flow. We complement LSCI data with Lomb-Scargle power spectrum analysis of capillary diameter dynamics obtained using two-photon microscopy. We apply the combined techniques to assess cerebral perfusion and pulsatility across nearly the entire lifespan of awake wild-type (C57BL/6) mice - from 18 to 81 weeks. Furthermore, at every time point, we perform additional measurements while anaesthetising animals using isoflurane or ketamine-xylazine, the most commonly used anaesthesia types. Our findings reveal the absence of an age-related decrease in perfusion, demonstrating that blood flow index and diameters remain remarkably stable over animal life. Pulsatility characteristics are consistent with the perfusion and show no change until the final weeks of observation, where an increase in arterial pulsation appears. In contrast, measurements taken under anaesthesia display a range of age-dependent and age-independent changes. Most strikingly, despite their distinct mechanisms and effects on the vasculature, isoflurane and ketamine-xylazine significantly increase perfusion pulsatility and affect diameter pulsatility. It highlights a previously overlooked detrimental effect of anaesthesia when assessing brain function but also suggests a new possible approach to control the blood-brain barrier permeability.

## 2 Results

### 2.1 Pulsatility and perfusion in the ageing brain

Microvascular perfusion and pulsatility measurements in ageing awake mice are presented in Fig. 1. Surprisingly, *BFI* and *PI_BF I_* do not change across the entire observation period, with variation in average values being below 5% for *BFI* and 5% for *PI_BF I_*between age groups (Fig. 1, A), highlighting the stability of the perfusion as opposed to age-related perfusion decline hypothesized in some studies. The average intra-group variation within the same timepoint is also low, 10% for *BFI* and 10% for *PI_BF I_*, which reflects the robustness of measurements and small physiological variability between animals. Average *BFI* and *PI_BF I_* images, as well as statistics on the number of identified segments, are shown in Supplementary Figures A1, A2 and A3, respectively. Notably, *PI_BF I_* of *≈* 0.17 in small arteries, *≈* 0.14 in the parenchyma, and *≈* 0.12 in small veins (Fig. 1, C) is not only age but also system-independent, which makes it a reference for healthy mice within the given age and vessel diameter. *D* and *PI_D_* measurements, shown in Fig. 1, B and D, follow almost the same pattern, except for the last time points, where the average venular diameter appears to increase after week 73. Interestingly, the average arterial diameter does not increase at the same time. Instead, arterial diameter pulsatility *PI_D_* is significantly increased from 0.09 at week 65 to 0.14 at week 81. Such observation could imply that arteries, contrary to the expectations, became more elastic rather than stiffer. Two-photon microscopy measurements show no significant changes in capillary flux, density, or diameter (Fig. 1, E-G), matching well with stable BFI throughout the observation period. The average capillary diameter also does not show a significant difference (Fig. 2, D), although the median diameter has increased at 83, matching with the venular diameter increase observed with LSCI. The signal-to-noise ratio of capillary pulsatility, estimated from the Lomb-Scargle periodogram, however, does show a significant decrease when comparing animals of 83 and 38 weeks.

**Fig. 1.**
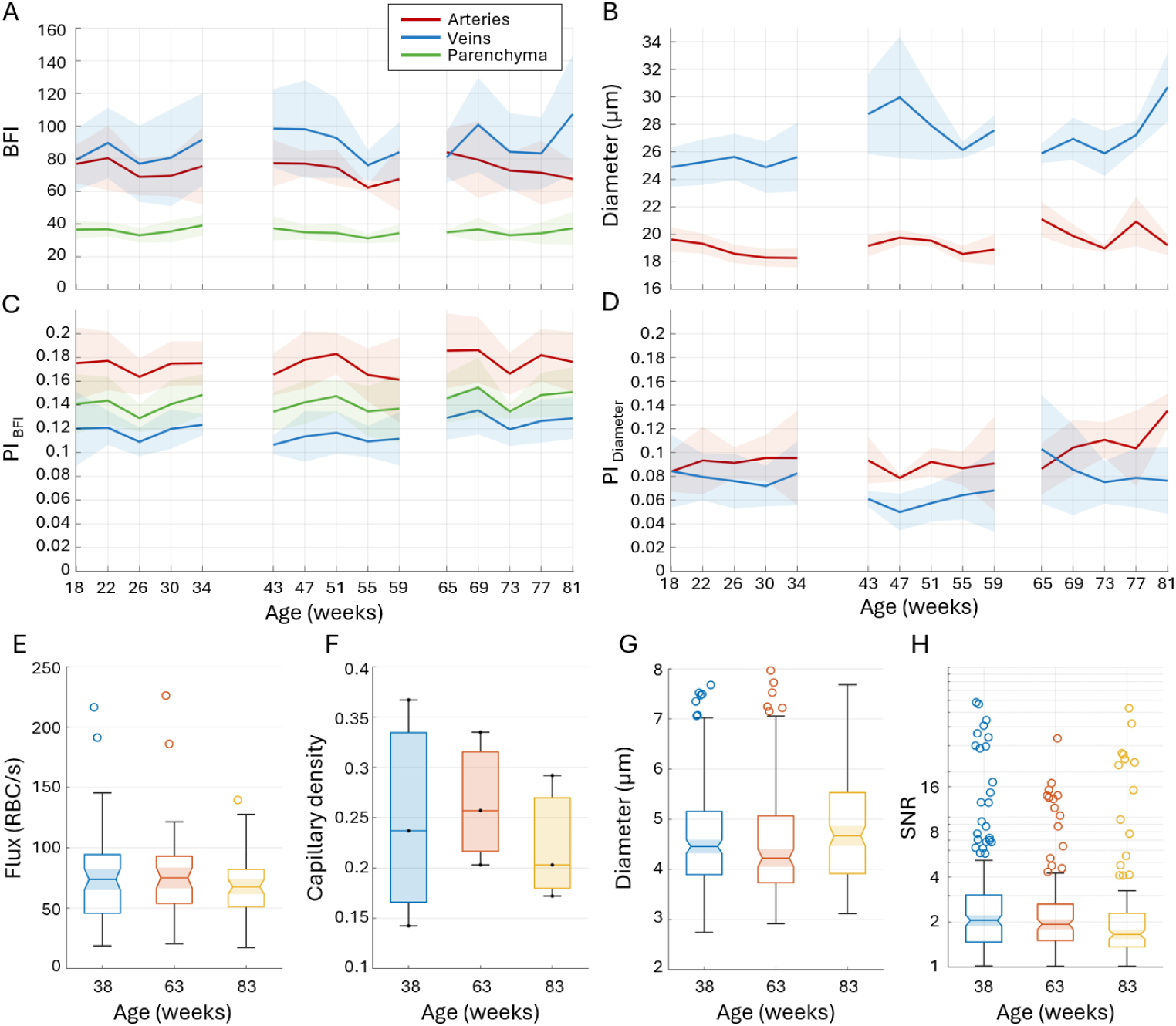
LSCI (A-D) and TPM (E-H) measurements of perfusion, diameter and respective pulsatility in ageing awake C57BL/6 mice. A - average blood flow index (*BFI*) in small arteries (red), veins (blue) and parenchyma (green) corresponding to the mouse age. B - average diameters (*D*) of respective arteries and veins. C - average BFI pulsatility index (*PI_BF I_*) for the respective vessels and parenchyma. D - average diameter pulsatility index (*PI_D_*) for the respective arteries and veins. Solid lines and shaded areas reflect the mean and standard deviation of the corresponding measurements across animals. Details on the number of animals and vascular segments for each measurement are shown in Supplementary Figure A1. E-H capillary flux, density, diameter and pulsatility signal-to-noise ratio estimated from TPM data.

**Fig. 2.**
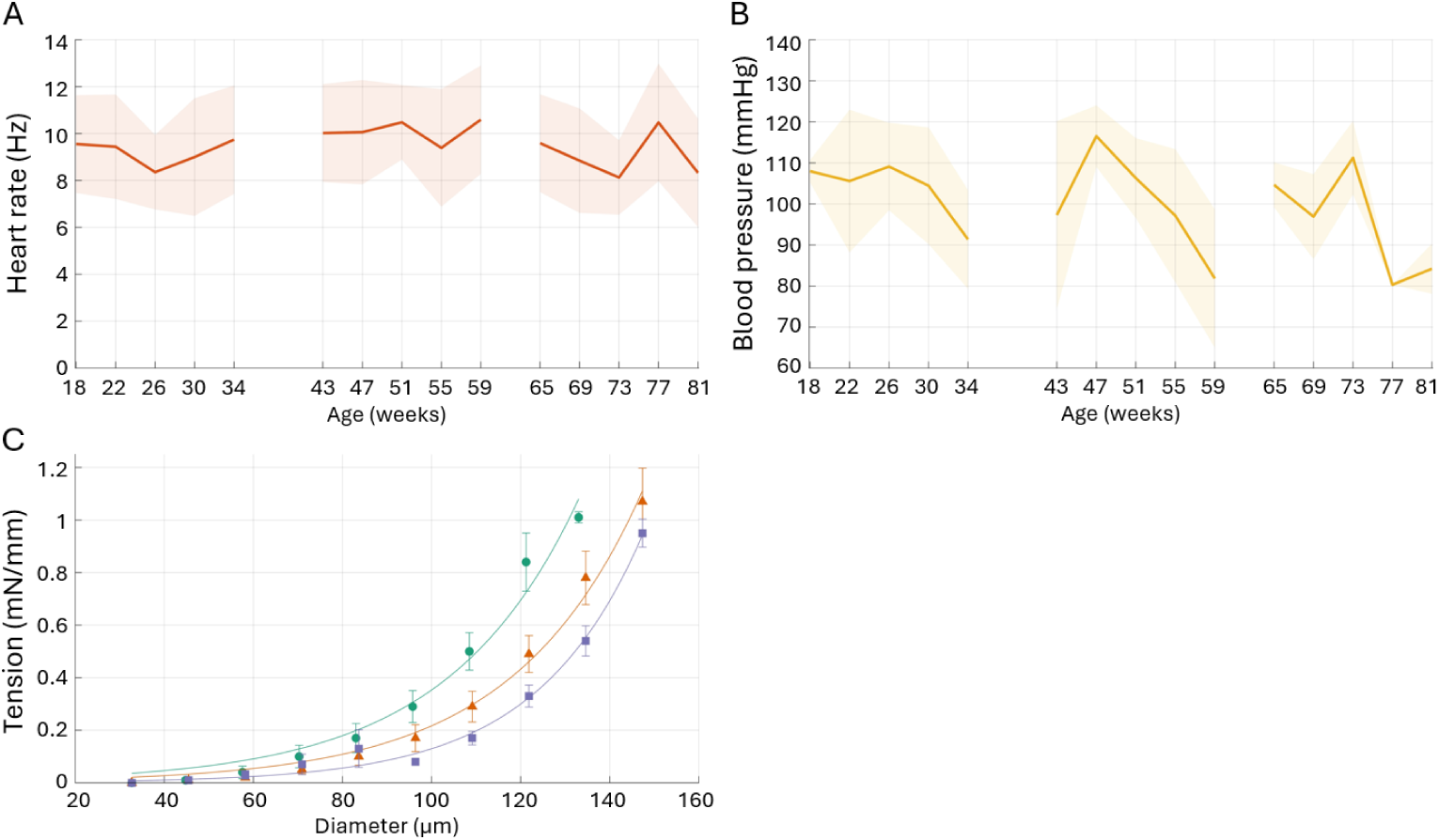
Systemic measurements (A,B) in ageing awake mice, as well as ex-vivo data (C, D). A - average heart rate measured from the LSCI data, corresponding to the data shown in Fig. 1. B - average blood pressure from the tail-cuff measurements at the end of each LSCI recording. Note that zero standard deviation in week 77 was caused by the system malfunction and, therefore, just one valid measurement. C - wall-tension curves from isometric wire myography measurements in isolated middle cerebral arteries 38, 63 and 83 weeks old for the green orange and purple lines respectively.

Systemic and ex-vivo measurements align well with the perfusion and pulsatility imaging. Average heart rate remained within 8.1-10.6 Hz, with less than 10% variation between the average for the age groups and less than 20% intra-group variation (Fig. 2, A). Similarly, the average blood pressure has generally remained between 80.3 and 116.5 mmHg with respective variations of *<* 10% and *<* 20% (Fig. 2, B). Neither heart rate nor blood pressure changes could be associated with the increased venular diameter and arterial pulsatility observed in the last time points (Fig. 1, B, D). The tension curves obtained with isometric wire-myography confirm the late-onset diameter pulsatility changes, as they show tension reduction with age (Fig. 2, E), implying that middle-cerebral arteries became more elastic.

### 2.2 Pulsatility along the microvascular network

Figure 3 shows how pulsatility and perfusion change along the microvascular network - from arteries to veins across all ages. Naturally, the blood flow index displays an inverse bell shape, as it first reduces with decreasing diameter on the arterial side, reaching a minimum in the parenchyma and increasing along with the diameter on the venular side (Fig 3, A-C). The BFI pulsatility index is consistently decreasing, from as high as 0.21 in large arteries to as low as 0.09 in large veins, fitting the flux pulsatility decrease shown in [24]. It reflects how flow oscillations are dampened along the elastic vascular network. The diameter pulsatility displays behaviour inverted compared to the BFI - large arteries and veins appear as non-pulsating (*PI_D_ <* 0.06), while the smallest vessel experiences the most extensive changes in the diameters (*PI_D_ >* 0.11). Furthermore, while the BFI pulse shape is consistently changing along the network, with the peak getting more Gaussian-like and more delayed (Fig 3, B), the diameter pulse shape mostly remains the same (Fig 3, C). More details on the exact correlations between different LSCI measurements can be found in the supplementary Figure A4.

**Fig. 3.**
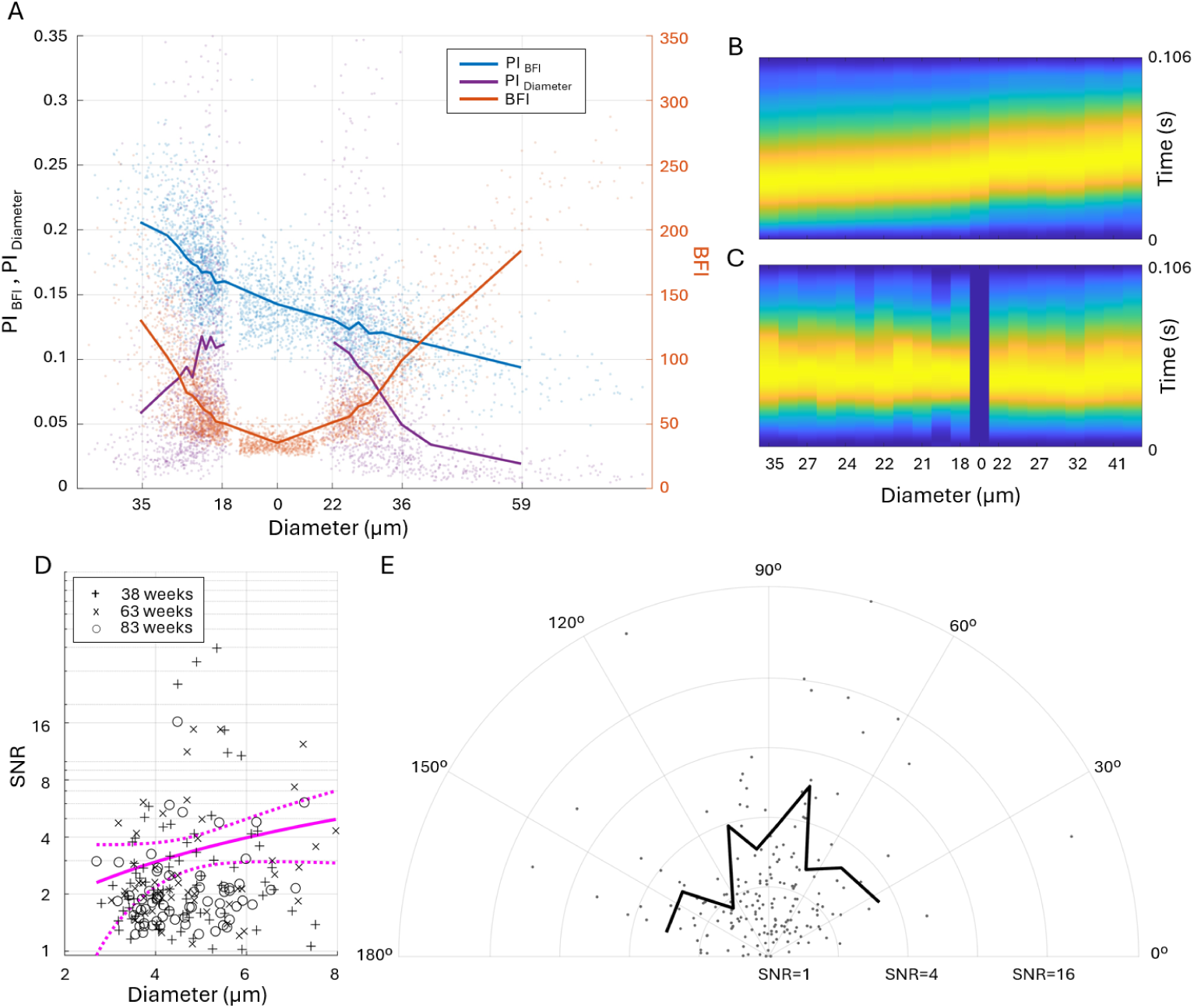
Change in pulsatility and perfusion across the microvascular network for all measurements. A – average diameter and perfusion pulsatilities *PI_D_* and *PI_BF I_* (yellow and blue lines, left Y-axis), and blood flow index *BFI* (red, right Y-axis) plotted as a function of the average diameter *D* across the imaged microvascular network. Markers indicate individual measurements. Note that parenchymal measurements, by definition, do not have a diameter value associated with them - the respective markers were assigned to a random diameter that is below the smallest vessel diameter for better visualisation. B - the shape of the BFI pulse, normalised for each average diameter and presented as a colour-coded map with time and diameter axis. C - the colour-coded map of the shape of the diameter pulse for different average vessel diameters. D - log-scale signal-to-noise ratio at the cardiac frequency calculated from Lomb-Scargle power spectrums of capillary diameters. Different marker symbols correspond to the measurements done at 38 (”+”), 63 (”x”) and 83 (”o”) weeks respectively. Magenta lines show the linear regression model and 95% confidence intervals calculated across all measurements. E - polar plot of capillary pulsatility SNR for different vessels orientation relative to the coronal plane.

Lomb-Scargle power spectrum analysis of the TPM data confirms the presence of the cardiac-cycle associated diameter fluctuations in capillaries (3 E, F). Linear regression of the signal-to-noise ratio at the cardiac frequency shows that pulsatility increases with the diameter, similar to changes in the absolute pulse magnitude for small vessels measured with LSCI. Interestingly, both LSCI and TPM data show severe pulsatility heterogeneity, where the diameter oscillations in a minority of vessels are several times stronger than for the majority of them (3 A, D). Furthermore, as shown in Fig 3, E the SNR is not correlated with the vessel orientation, confirming that it is associated with vascular pressure wave propagation and not with the displacement of the entire brain. Such displacement is characterised in Figure 4. The estimated SNR for vessel displacement is much higher than for capillary pulsatility and, unlike the latter, is strongly orientation-dependent (4 A, B). The displacement is most pronounced in vessels oriented at roughly 40 degrees to the coronal plane, which matches the expected orthogonal direction to the pressure wave travelling through the brain tissue from the spinal artery. Most interestingly, the displacement SNR also reduces with age, possibly reflecting changes in tissue rigidity.

**Fig. 4.**
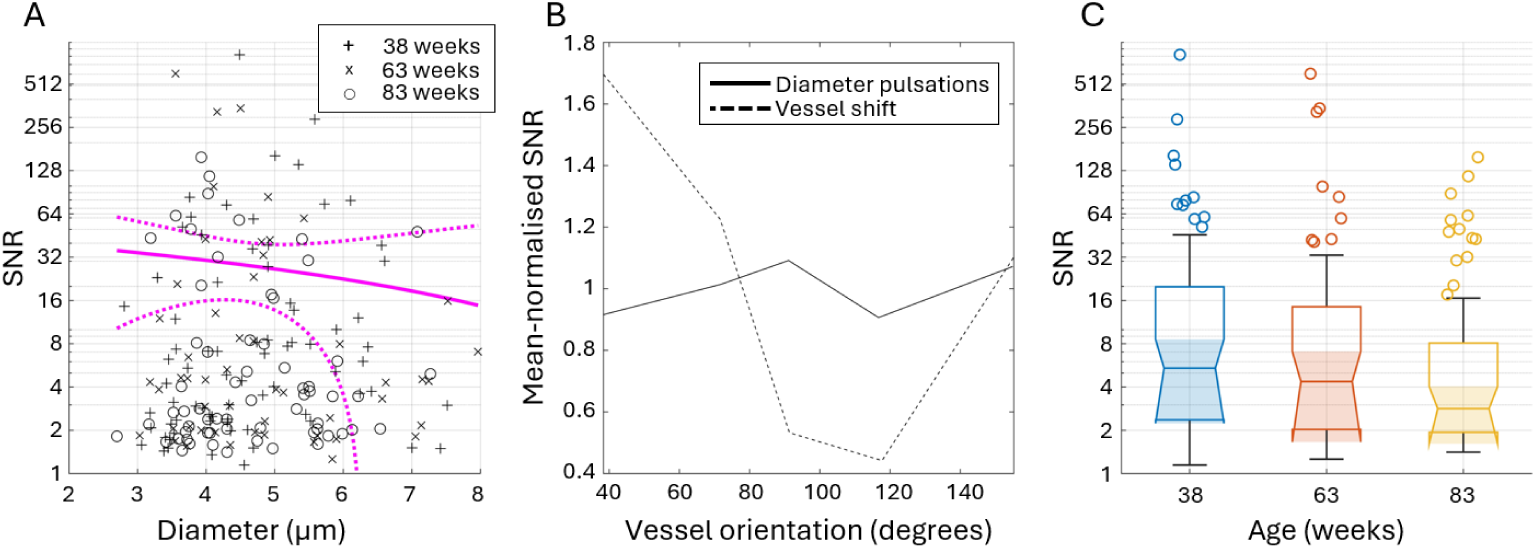
Lomb-Scargle analysis of capillary wall dynamics associated with the displacement of the entire brain during cardiac cycle. A - log-scale signal-to-noise ratio at the cardiac frequency calculated from Lomb-Scargle power spectrums for capillary wall coordinates change. Different marker symbols correspond to the measurements done at 38 (”+”), 63 (”x”) and 83 (”o”) weeks respectively. Magenta lines show the linear regression model and 95% confidence intervals calculated across all measurements. B - mean-normalised SNR of displacement and capillary pulsatility for different vessel orientations. Capillaries orientation is grouped in 20*^◦^*-wide bins. C - displacement SNR grouped by the animals’ age.

### 2.3 Confounding effects of isoflurane

Blood velocity and vessel diameter are known to be strongly affected by isoflurane due to its potent vasodilatory action. Figure 5 further expands our understanding of such effects, providing insight into perfusion and pulsatility response to low-dose isoflurane compared to the awake condition in ageing mice. As expected, BFI and diameters have significantly increased but, interestingly, in an age-dependent manner - the increase is most substantial in 18 weeks-old animals, upward of 200% for arterial BFI and 34% for diameters, and goes down to *≈* 100% and *≈* 20% respectively by the age of 43 weeks (Fig 5, A, B). Perfusion pulsatility index *PI_BF I_* pulsatility index has also significantly increased, in line with the BFI changes (Fig 5, C). Intriguingly, pulsatility seems to increase relatively less in arteries and the most in veins - for the youngest age group, the average *rPI_BF I_* is 1.7 in arteries, 1.8 in the parenchyma and 1.9 in veins. The tendency is the same, albeit less pronounced, for the second and the third age groups with the respective average *rPI_BF I_* of 1.5 and 1.42 in arteries, 1.51 and 1.44 in the parenchyma and 1.6 and 1.47 in veins. The diameter pulsatility index (Fig 5, D), on the other hand, has only significantly increased in veins, with average *rPI_D_* being *≈* 1.95, 1.7 and 1.4 in the respective age groups. Arterial *PI_D_* almost did not increase compared to the awake measurements despite the increased flow pulsatility. It has only increased by *≈* 14% for the first age group, increased by *≈* 4% in the second group and has even decreased by *≈* 5% in the oldest group. Heart rate and blood pressure (Fig. 5 E, F) have both decreased by 20 *−* 25% compared to the awake condition but did not display age-dependent dynamics. More details on how the isoflurane affects the pulsatility along the microvascular network are shown in Supplementary Figure A6, A,C,D.

**Fig. 5.**
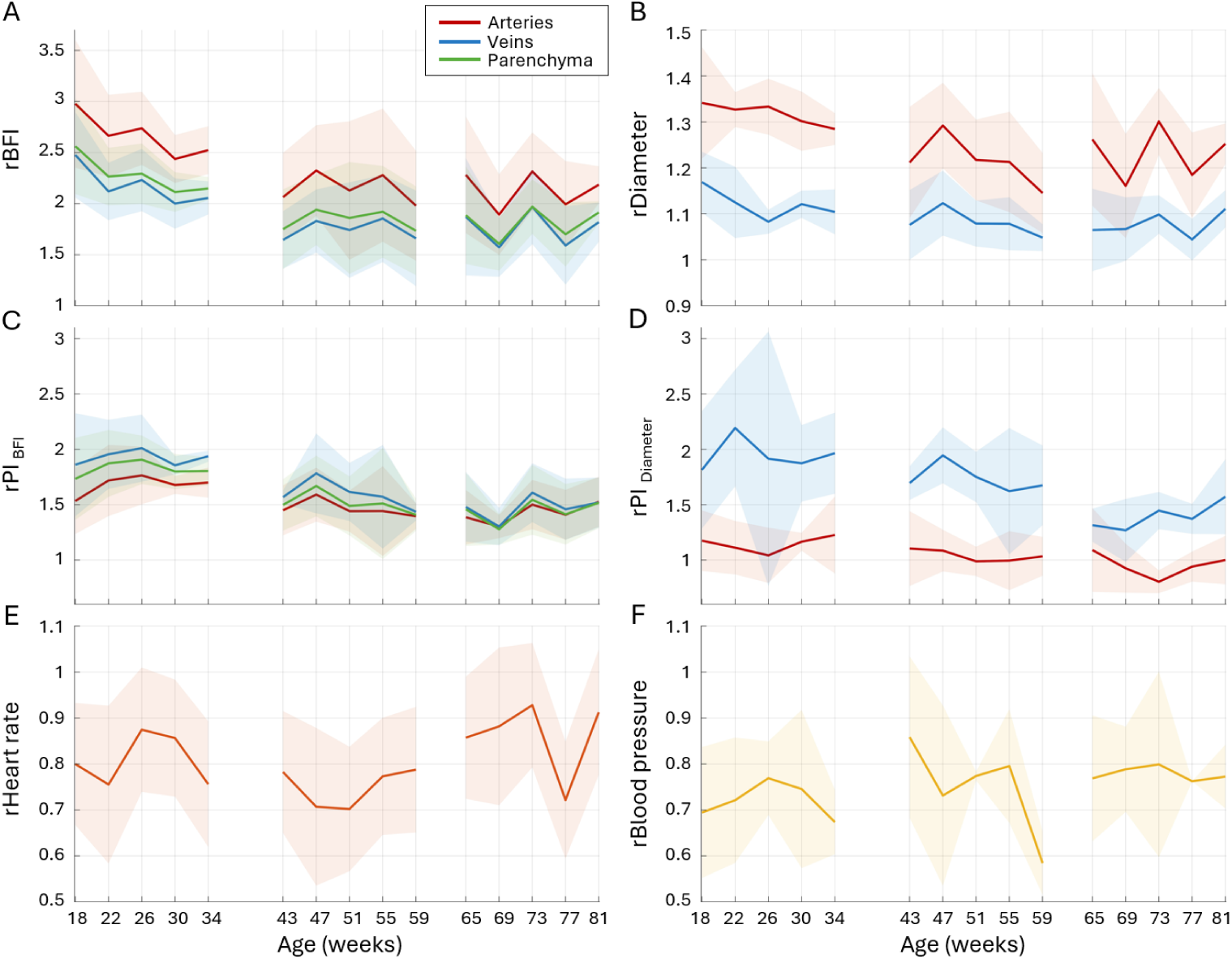
Age-dependent effects of isoflurane on perfusion and pulsatility. The results are displayed as a relative change compared to the measurements taken in awake animals. A - average relative blood flow index change (*rBFI*) in small arteries (red), veins (blue) and parenchyma (green), respectively. B - average relative diameter (*rD*). C - average relative BFI pulsatility index change (*rPI_BF I_*). D - average relative diameter pulsatility index change (*rPI_D_*). E - average relative heart rate change. F - average relative blood pressure change.

### 2.4 Confounding effects of ketamine-xylazine

Unlike isoflurane, the ketamine-xylazine combination is not a vasodilator and has almost no effect on average diameters (less than 5% increase in arteries, Fig 5, B). Nevertheless, perfusion has increased by 10-40% (Fig 5, A), but without a clear age-associated pattern. Despite a much weaker effect on perfusion and diameter, ketamine-xylazine has caused an even more profound increase in pulsatility compared to isoflurane. Under ketamine-xylazine, the BFI pulsatility index has, on average, increased by *≈* 40 *−* 80% in arteries, *≈* 50 *−* 90% in the parenchyma and *≈* 75 *−* 120% in veins. Diameter pulsatility has also risen, with average *PI_D_* increasing by 50-160% in both arteries and veins. Such dramatic changes in pulsatility are accompanied by a two-fold decrease in the heart rate (Fig 5, E) and an absence of changes in blood pressure. The latter remained primarily similar to the awake condition, alas, appears to decrease *≈* 20% in the youngest age group. More details on how the ketamine-xylazine combination affects the pulsatility along the microvascular network are shown in Supplementary Figure A6, B,E,F.

## 3 Discussion

In the present study, we suggest a new, highly robust approach for simultaneous perfusion and diameter pulsatility characterisation using full-field blood flow imaging and two-photon microscopy and use it to advance the understanding of pulsatility in the brain cortex of ageing awake and anaesthetised mice. Figure 1, A, shows how remarkably stable the cerebral perfusion of wild-type mice is. It contributes to the growing body of studies suggesting that there is no age-related perfusion decrease [28, 29] in the cortex, at least in the awake wild-type mice younger than 81 weeks. While there are reports supporting the opposite [30–32], we hypothesise that it is likely a consequence of different animal preparation approaches and the use of anaesthesia, which we will discuss below. The perfusion stability we observed is supported by the absence of age-dependent changes in the heart rate, blood pressure, capillary flux, density and diameters. We show that microvascular BFI pulsatility is similarly stable and also that it is comparable in magnitude with hyperaemic responses observed during functional stimulation [33]. From week 18 to week 81, the average *PI_BF I_* maintains the value of *≈* 0.18 in small arteries, *≈* 0.14 in parenchyma and *≈* 0.12 in small veins (1, C). Such stability can be seen as either an absence of vascular or hemodynamic changes or a complex combination of changes that balance each other, keeping the cerebral perfusion stable. The latter seems more likely due to confirmed functional vascular and neurovascular abnormalities in the ageing mouse brain [33, 34] that occur despite stable perfusion. The BFI pulsations persist throughout the entire microvascular network - starting from an average magnitude of 21% in the first branches of middle cerebral arteries with the diameter of *≈* 35*µm* and reaching as low as 9% in the largest segmented veins with the diameter of *≈* 59*µm* (see Fig. 3). It is natural to assume that the pulsatility values and the shape of the *PI_BF I_* decline will change in conditions associated with microvascular abnormalities, as we see in animals under anaesthesia (Supplementary Figure A6). More specifically, a higher *PI_BF I_* and the slowdown of its decrease can be associated with increased arterial stiffness or hemodynamic resistance (e.g. due to capillary rarefaction). A decrease in *PI_BF I_* and, possibly, faster decline can be associated with upstream occlusion or flow reduction (e.g. in stroke or transient ischemia).

The diameter pulsatility aligns with the *BFI* stability until week 73. After that point, the average vein diameter increases significantly, more than 20% compared to the first timepoint of the same age group. It aligns with a non-significant but noticeable increase in the median capillary diameter and a decrease in capillary pulsatility (Fig. 1, G, H), as measured from the TPM line scans. Surprisingly, this change is not reflected in arterial diameter or *BFI* in arteries or parenchyma. Instead, average arterial diameter pulsatility *PI_D_* during week 81 becomes significantly higher than in week 65, increasing from 0.09 to 0.14 (Fig. 1, D). It can imply either a stronger pulse reaching the brain, arteries becoming more elastic at this age or both. A stronger pulse to reach the brain would require either an increased cardiac output, which is unlikely, or increased stiffness of large arteries, e.g., aorta. While little is known about vascular stiffness in ageing mice, there are reports of a mild increase in aorta stiffness by the age of 80 weeks [16, 35, 36] under anaesthesia. Contrary to the increase in aortic stiffness, our findings from isometric wire myography show a decreased tension and, therefore, higher elasticity in middle cerebral arteries with age (Fig. 2, D). Initially counterintuitive, decreased stiffness, and therefore higher diameter pulsatility in the small arteries (Fig. 1, D), might be a mechanism that counters increased pulsation caused by stiffer large vessels and therefore protects the brain’s capillaries. Aside from the changes in the late age, the distribution of average *PI_D_* across the microvascular network, shown in Figure 3, is particularly interesting. It starts low (*PI_D_ <* 0.05) in arteries larger than 30*µm* in diameter and then increases for smaller vessels, reaching the peak of *PI_D_ >* 0.1 in smallest resolvable arterioles and venules with *≈* 12*µm* diameter, and then decreasing to its lowest in larger veins. Together with the rapid decrease in the *PI_BF I_* in arteries below *≈* 12*µm* diameter, it suggests that these small vessels play a prominent role in microvascular pulse dissipation and are likely candidates for the “drivers” of pulse-associated fluid motion [1]. Crucially, the diameter pulsatility appears to be strongly heterogeneous, with both LSCI and TPM data showing that at the same average diameter, some vessels pulsate several times more than others. Understanding the cause of heterogeneity requires further exploration, but we assume that it can be related to the tissue’s structural properties in the proximity of the vessels or the local tone of mural cells, including pericytes. At the same time, it highlights that some of the small vessels might be especially vulnerable to abnormal increases in pulsatility, which are likely to disrupt the blood-brain barrier.

Following up on the observations above, we acknowledge that the pulsatility and, possibly, perfusion may change more in the later age, closer to the 105-120 weeks life expectancy of the C57BL/6 male mice [37]. In such perspective, 81 weeks “correspond” to only 60 human years, at which cerebral perfusion decline may not be present [38]. At this age, mice also show almost no cognitive decline [39], similar to humans where dementia symptoms before 65 years are uncommon and referred to as “early-onset” dementia [40]. Intriguingly, however, assuming perfusion and cognition change in older animals suggests that the diameter pulsatility changes we observed at week 81 might precede other abnormalities, making their further exploration pivotal for understanding age-related brain perfusion changes and associated pathologies.

General anaesthesia, although known to have multiple systemic and local effects, is still broadly applied in research - either in settings where an awake animal preparation is impossible or to save the effort required for it. Isoflurane and ketamine-xylazine are, in particular, among the most common anaesthesia choices [41]. Isoflurane is a general inhalation anaesthetic which conveniently allows long-term stability when administered via mask or intubation. As an agonist of adenosine triphosphate-sensitive K+ channels, however, it causes hyperpolarisation of the smooth muscle cells and, therefore, potent vasodilation [42, 43], raising concerns for interpretation of the perfusion measurements and functional responses. Our results show that even a “minimal” dose of isoflurane can cause up to 200% in BFI and 34% in arterial diameter in young animals (Fig. 5, A, B). We quote “minimal” here since we used the lowest isoflurane concentration, which allowed stable cerebral blood flow in our experimental setup (see Methods for details). Most critically, we show that the isoflurane-associated increase in perfusion and diameters strongly weakens with age (by *≈* 60%), as was indirectly reported in earlier studies [44]. Such age-dependent effects would make any ageing-related results challenging to interpret, especially considering that heart rate and blood pressure remain age-independent (Fig. 5, E, F). It might also be one of the key reasons some studies have previously reported a decline of cerebral blood flow in ageing mice [30–32]. The ketamine-xylazine combination, unlike isoflurane, allows for simple injection routes, does not require gas induction equipment, and has less pronounced effects on the brain vasculature and perfusion. The latter is also supported by our results (Fig. 6, A, B), which show nearly no changes in vessel diameter and, at the most, a 10% increase in blood flow under ketamine-xylazine anaesthesia compared to the awake animals. Nevertheless, both ketamine and xylazine can cause systemic changes which will inadvertently affect brain perfusion. Ketamine has a sympathomimetic effect where it increases heart rate and blood pressure, while xylazine causes cardiovascular depression, including bradycardia, hypotension, and decreased cardiac output. The systemic effects, especially from slower and longer-lasting xylazine, are clearly reflected in our heart rate measurements (Fig. 6, A, B), displaying a 50% reduction compared to the awake condition. Unlike isoflurane, however, ketamine-xylazine has shown no explicit age-dependent perfusion dynamics, possibly making it more optimal for ageing studies requiring anaesthesia.

**Fig. 6.**
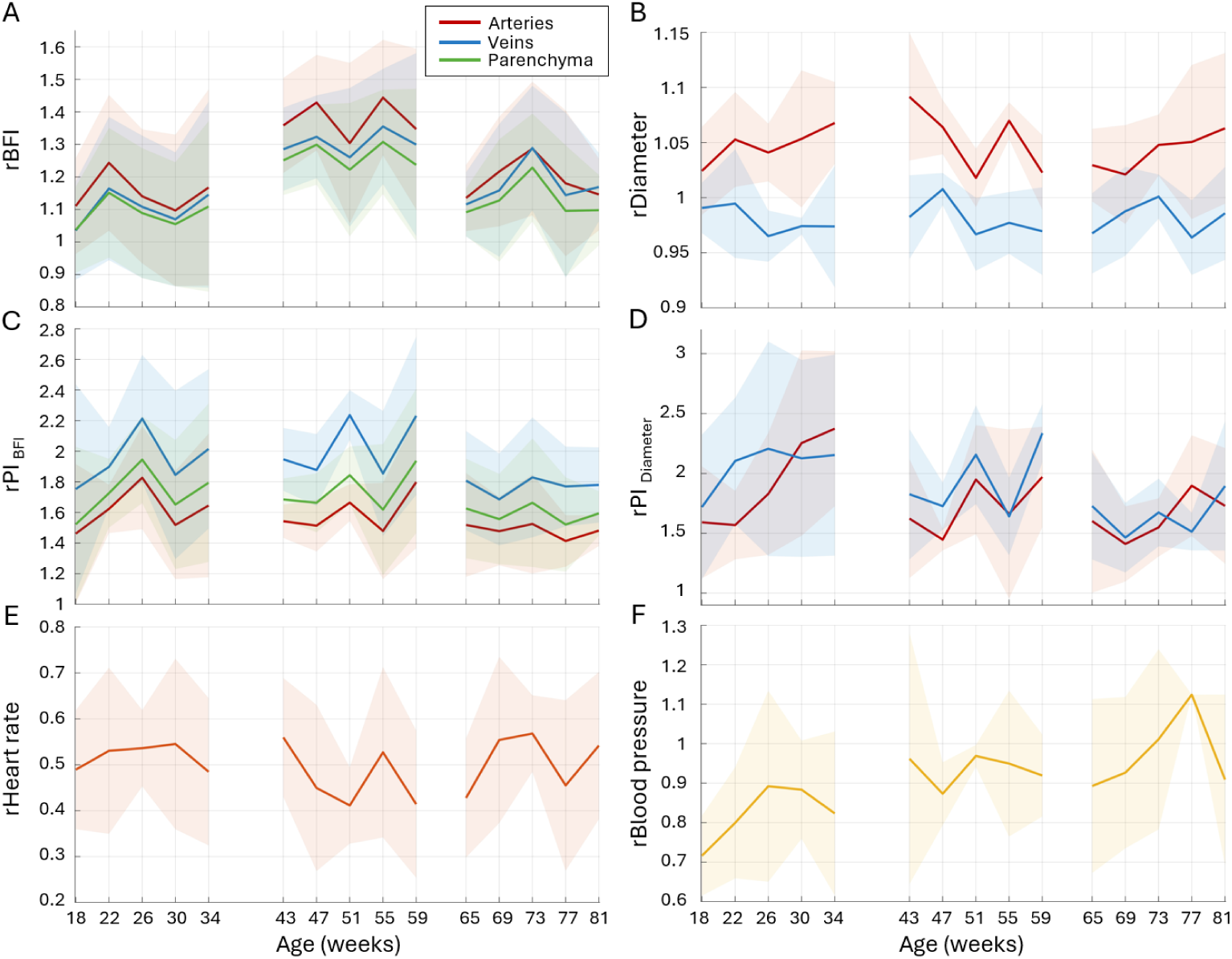
Effects of ketamine-xylazine anaesthesia on perfusion and pulsatility. The results are displayed as a relative change compared to the measurements taken in awake animals. A - average relative blood flow index change (*rBFI*) in small arteries (red), veins (blue) and parenchyma (green), respectively. B - average relative diameter (*rD*). C - average relative BFI pulsatility index change (*rPI_BF I_*). D - average relative diameter pulsatility index change (*rPI_D_*). E - average relative heart rate change. F - average relative blood pressure change.

We have also addressed how anaesthesia affects microvascular pulsatility in the brain cortex. The perfusion pulsatility index *PI_BF I_* has significantly increased for both isoflurane and ketamine-xylazine despite their distinct effects on the perfusion. As Figure 5, C, shows, in the isoflurane case, it follows the age-dependent dynamics. *PI_BF I_* rises the most in the youngest age group and less in the older animals, with the respective increase of 50 *−* 100% and 40 *−* 70% depending on the vessel type. For the ketamine-xylazine (Fig. 6, C), despite a lower *BFI* increase compared to isoflurane, *PI_BF I_* has increased even more, showing a 50 *−* 120% increase in all age groups. Notably, the effect differs depending on the vessel type - the *PI_BF I_* increased relatively less in arteries, more in the parenchyma and the most in veins. It may reflect arteries reaching the capacity limit to compensate for perfusion fluctuations, allowing a disproportionately stronger pulse to reach capillaries and veins. Such behaviour can occur in pathologies where the vascular network’s pulse dissipation ability is compromised, e.g. due to increased stiffness or a stronger pulsation. Counterintuitively, it is not necessarily associated with vasoconstriction but can also be caused by extensive vasodilation, as confirmed by the diameter pulsatility changes under anaesthesia. For isoflurane, only venular *PI_D_* is increased (by 40 *−* 110% depending on age, Fig. 5, D), while for ketamine-xylazine both arterial and venular *PI_D_* is nearly doubled (Fig. 6, D). We suggest a straightforward explanation of such dynamics. Isoflurane causes potent dilation of arteries, making them lose the capacity to change the diameter and dissipate the pulse energy. Thus, the increased pulsatile flow, reflected in higher *BFI* and *PI_BF I_*, propagates through the arterial side and reaches veins at levels comparable to arterial pulsatility in healthy awake animals, causing strong diameter pulsations. Ketamine-xylazine does not dilate arteries, but the heart rate is reduced to half of its value in awake animals while the *BFI* is maintained. That is only possible if every heartbeat becomes stronger, causing increased pulsatility. Unlike the isoflurane case, however, the arteries are not dilated and, thus, to some degree, can compensate for the perfusion fluctuations with passive diameter changes, which reflects not only in arterial *PI_D_* but also *PI_BF I_*. For ketamine-xylazine, a relative increase in parenchymal pulsatility is closer to the veins, while for isoflurane, where arteries have nearly lost their compensatory ability, change in parenchymal pulsatility is closer to the venular side (C panel of Figures 5 and 6). Nevertheless, even for ketamine-xylazine, arteries seem to reach the limit of how much they can compensate for perfusion fluctuations, as the pulsation strongly propagates to the venular side. Overall, we found that both isoflurane and ketamine-xylazine significantly amplify pulsatility in the brain. Consequently, it can enhance waste clearance by augmenting pulse-associated vascular motion, disrupt blood-brain barrier integrity by simultaneously dilating vessels and increasing intravascular pressure fluctuations, and alter functional responses, for example, by activating Piezo1 receptors [45, 46]). These prospects highlight the importance of pulsatility effects when interpreting the results obtained under anaesthesia but also suggest a potential pathway for anaesthesia to improve drug delivery across the blood-brain barrier that is unrelated to tight junction proteins [47].

The obtained results also validate the robustness of the proposed method. The optimized design of the LSCI system we developed (see Methods for details) allowed measurements with less than 10% intra-group variation and less than 10% intra-animal variation across all time points. Considering physiological variability in perfusion, which undeniably contributed to the measurement variation, it confirms that the system provides reliable absolute BFI measurements and excels in longitudinal studies or studies comparing perfusion in different animal models. It can be easily implemented as a part of other imaging systems, e.g., multi-photon microscopy or optical coherence tomography, and it provides unparalleled full-field, high frame-rate insight into vascular and perfusion dynamics. Moreover, the pulsatility measurements are unitless and system-independent and thus can be compared directly between studies, allowing the use of presented *PI_BF I_* and *PI_D_* as a reference in healthy ageing wild-type animals. The TPM and ex-vivo data align well with LSCI, supporting our observations. Nevertheless, we must acknowledge several limitations and considerations for future studies. First and foremost, despite becoming the golden standard for optical brain imaging, chronically installed cranial windows might affect the tissue underneath and the brain as a whole [48]. Therefore, further research is required to understand how chronic cranial windows might alter vascular stiffness, pulsatility and perfusion and whether the interpretation of our results has to be changed accordingly. From the imaging perspective, a larger field of view and higher optical resolution would improve the accuracy and precision of the results. We deliberately limited the field of view to 1024×512 pixels as a trade-off to reduce the data volume. When not limited by the storage capacity, the field of view can be increased by up to 8 times (2048×2048 pixels) for the same consumer-grade CMOS sensor while keeping *≈* 200 Hz framerate. In such a case, by changing the objective, one could record more vessels at a higher resolution, getting more reliable data in small vessels. The latter is directly related to limitations of the segmentation algorithm as the diameter estimation error will naturally increase for pixel-wise smaller vessels [49–52]. Furthermore, a question might arise regarding inducing ketamine-xylazine following the isoflurane in our experimental protocol, which we addressed by performing pilot experiments. Still, we must acknowledge that a more thorough independent investigation of ketamine and xylazine actions would be needed to understand their effect on microvascular pulsatility. Despite the abovementioned limitations, we see the presented results and methods seminal to future research in microvascular pulsatility and understanding its role in brain function.

## 4 Conclusion

In conclusion, we have presented a new approach for simultaneous full-field characterisation of microvascular pulsatility and, in combination with two-photon microscopy and ex-vivo analysis, applied it to advance the current understanding of perfusion and pulsatility in the brain cortex of awake and anaesthetised ageing mice. The average *BFI* variation remained below 10% for measurements in each animal, within each group and between groups, showing the absence of the age-related decline from 18 to 81 weeks old and confirming the method’s robustness. Independent of the age, the BFI pulsations declined along the microvascular network, from 0.21 to below 0.08, with roughly 37% of the decrease happening in small arteries with the average diameter of 12-23 *µ*m. Such magnitude of the BFI change is comparable to peak hyperaemic response during functional activation, highlighting that the pulsatility role in brain perfusion should not be underestimated. At the same time, the diameter pulsatility analysis implies that arteries, counter-intuitively, become more compliant with age. Average arterial pulsatility has increased from 0.1 to 0.15 between weeks 65-81, accompanied by 20% increase in venular diameter and 10% increase in median capillary diameter. Isometric wire myography and histology confirm this observation, showing that older arteries develop weaker tone and contain less collagen. We argue that an increase in microvascular elasticity might act as a protective mechanism that responds to early increases in aorta stiffness and, therefore, a stronger pulse reaching the brain. It might also be a predecessor of later brain perfusion and pulsatility changes. Furthermore, we show that isoflurane and ketamine-xylazine cause a complex effect on pulsatility, which has to be accounted for when using these types of anaesthesia. Ketamine-xylazine caused an increase reaching more than 100% in perfusion and diameter pulsatilities across the entire microvascular network despite having little effect on the perfusion and average vessel diameters. A “minimal” dose of isoflurane has caused significant increases in *BFI*, perfusion pulsatility, diameters of both arteries and veins and venular diameter pulsatility. Most critically, the isoflurane effect appears to be strongly age-dependent, as the respective increases get lower with age, which is crucial for any ageing-related study.

## 5 Methods

### 5.1 Ethical approval declarations

All experimental protocols were approved by the Danish National Animals Experiments Inspectorate (permit no.: 2022-15-0201-01188) and conducted according to the respective guidelines and the Directive 2010/63/EU of the European Parliament on the protection of animals used for scientific purposes.

### 5.2 Animal preparation

In total, 15 male C57BL/6 mice (Janvier, Denmark), split into three age groups (each n=5) - starting at 18, 43 and 65 weeks were used for the study. Three animals, n=2 43 weeks old and n=1 65 weeks old, did not survive the complete protocol and, therefore, were excluded from the analysis. All animals were acclimatised in their cages for at least seven days and underwent surgical installation of a chronic cortical window over the left barrel cortex. For surgical procedures, animals were anaesthetised in an induction chamber with 3% isoflurane in 100% oxygen and reduced to 1.5-2% isoflurane once moved to a homeothermic heating pad to maintain core body temperature at 37*^◦^*C. Before surgery, analgesic (Buprenorphine, 0.1 mg/kg body weight), anti-inflammatory (Carprofen, 5mg/kg), antibiotic (Ampicillin, 200 mg/kg), and edema-reducing corticosteroid (Dexamethasone, 4.8 mg/kg) were administered intraperitoneally according to body weight. All fur was removed from the scalp with a rodent hair trimmer and Veet hair removal cream. Local anaesthesia (Xylocaine, 2mg/ml) was injected subcutaneously at the surgical site after removal of the hair. A small section of skin was removed from the skull, and the edges were secured to the bone using tissue adhesive (3M Vetbond). The periosteum was scraped off the exposed skull with a scalpel, and the skull was scored to increase surface area and thus decrease the risk of headbar detachment. A craniectomy was performed without damaging the dura, and the location was determined based on skull size and mouse age, but approximately 1.5–2 mm AP, 3 mm ML to Bregma. If any bleeds resulted from the craniectomy, these were stopped with a sterile saline-soaked hemostatic sponge before the placement of the glass window. The brain was ‘wiped’ clean with a saline-soaked hemostatic sponge to remove as many bone particles as possible to reduce bone regrowth. A small drop of Kwik-Sil adhesive (World Precision Instruments, Inc.) was placed on the exposed brain, and an optically transparent round glass coverslip Ø 4mm diameter was placed onto the window and fixed to the skull using Loctite light-cured glue. A metal head plate was glued to the skull to fix the head during imaging sessions. The entire skull was then covered with light-cured Relyx dental cement to seal the surgical site and fix the head plate.

After a successful surgery, the mouse was placed in a heated recovery chamber before returning to a new, clean cage. During the recovery period of a minimum of 10 days, weight and behaviour were closely monitored, and the mice received pain relief medication (Buprenorphine, Carprofen), antibiotic (Ampicillin), and fluids (saline) IP for four days following surgery as well as soft food at the bottom of the cage to encourage weight gain. Following the recovery period, the awake mice were habituated to restraint and trained in a mock Laser Speckle Contrast Imaging (LSCI) environment to limit motion artefacts and stress to obtain a stable blood flow during imaging. The training lasted for ten days, starting with 15 minutes of restraint and increasing by 15-minute increments daily until reaching two hours. The mice were rewarded with sweetened condensed milk during the habituation. At all times, animals were housed in 12 h light/dark cycle, 22-24*^◦^*C, 55 *±* 10% humidity, with *ad libitum* access to food and water.

### 5.3 Laser Speckle Contrast Imaging of microvascular pulsatility

#### 5.3.1 Experimental protocol

Before imaging, animals were lightly anaesthetised for 3 minutes in 3% isoflurane in 0.8L/min oxygen and moved to the imaging stage with a homeothermic heating pad to maintain a core body temperature of 37*^◦^*C. The mice were head-fixed to the imaging stage using clamps attached to the metal head plate. Once restrained in the imaging setup, a tail cuff (Kent Scientific) was placed on the mouse’s tail to allow blood pressure measuring, and the cuff was gently taped to the heating pad to keep it in place and the tail warm. The animals were allowed to recover from the initial isoflurane induction during the system setup and focusing for at least 15 minutes.

In each LSCI session, animals were imaged under three conditions while keeping the same field of view (Fig. 7, A): awake, isoflurane (1.5%) and ketamine-xylazine (75 mg/kg ketamine, 10 mg/kg xylazine). Respective blood pressure measurements were taken using a tail-cuff system at the end of each recording. Following the awake recording, a mask was placed over the mouse’s head, and the mouse was induced with 3% isoflurane and 0.8L/min oxygen for 3 minutes. Isoflurane was then decreased to 1.5%, which we found to be the lowest concentration at which stable blood flow can be achieved in our experimental setup (mask tightness and exhaust power). At lower values, *BFI* strongly fluctuated, most likely reflecting instabilities in anaesthesia condition, breathing and, therefore, isoflurane concentration in the bloodstream and its effect on blood vessels. LSCI recording was performed after the blood flow had stabilised but not less than after 20 minutes of anaesthesia. Isoflurane was shut down, and the mask was removed after completing the respective blood pressure measurement. After the mouse showed signs of recovery from anaesthesia (but not less than 15 minutes), a ketamine-xylazine cocktail was injected intraperitoneally. LSCI data was recorded 30 minutes after the injection. The procedure described above was repeated every four weeks for five months for each mouse. It is important to note that the 3-step protocol described above was chosen to minimise the number of imaging sessions and stress for the animals. Recovery after the isoflurane anaesthesia is typically within minutes [53], so it has always preceded ketamine-xylazine, whose effect lasts several hours, making randomised order experiment design unfeasible. We have also performed pilot trials without isoflurane to account for its possible impact on the ketamine-xylazine and found no significant differences in perfusion.

**Fig. 7.**
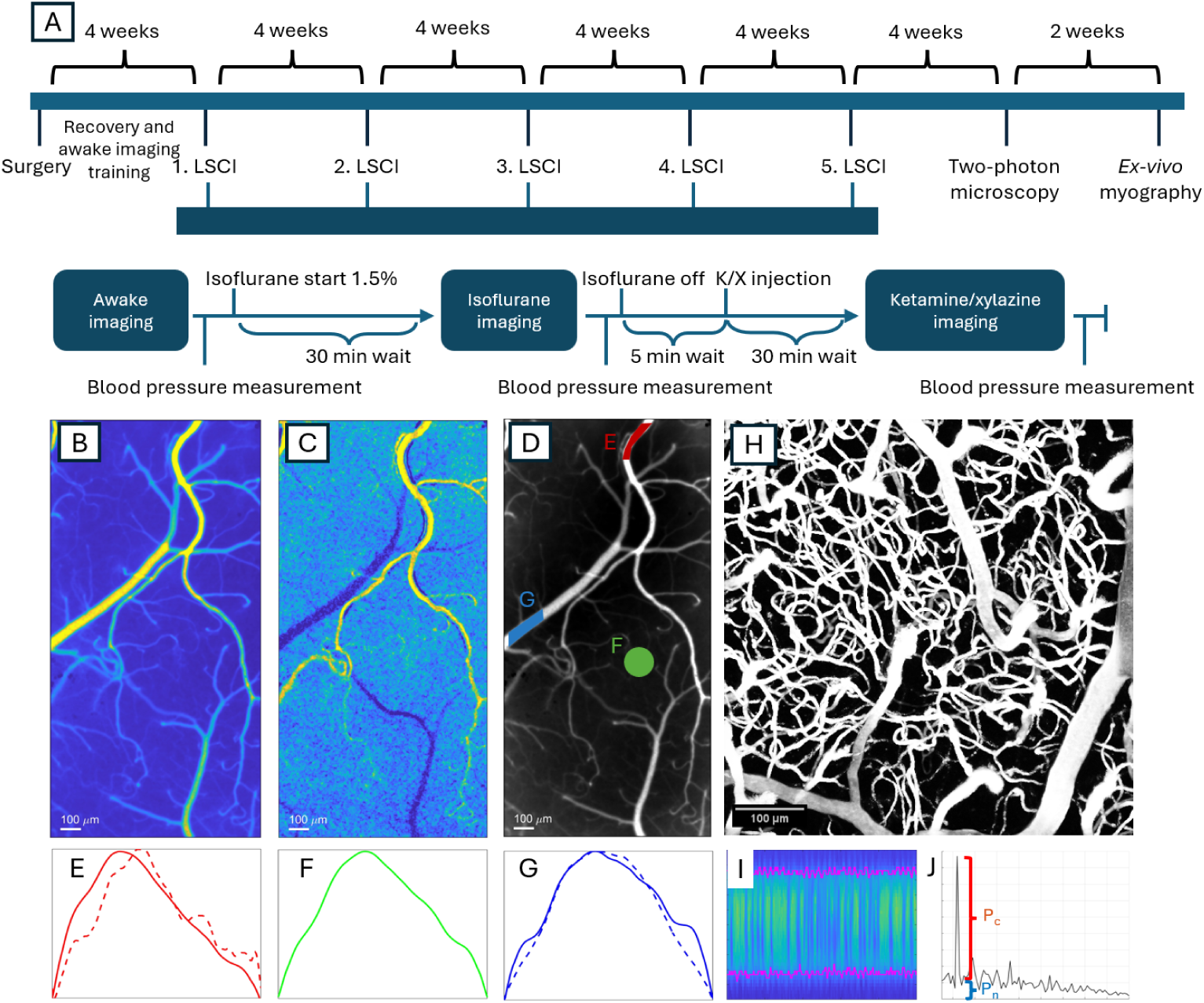
Experimental protocol and pulsatility imaging. A - timeline of procedures (top row) and the 3-step LSCI recording protocol (bottom row) for each mouse. B - exemplary average *BFI* image. C - exemplary *PI_BF I_* image of the same animal as in B. D - example of arterial (red), parenchyma (green) and venous (blue) ROIs placement. E, F, G - examples of the BFI (solid lines) and diameter (dashed lines) changes over the representative cardiac cycle in the corresponding ROI types in D. H - two-photon microscopy angiogram. I - an example of the crossectional capillary line scan is one in which the X-axis reflects the time and the Y-axis reflects the coordinates along the scan. The magenta colour denotes the detected vessel boundaries, which are then used for diameter estimation and consecutive power spectrum analysis. J - an example of the Lomb-Scargle periodogram of capillary diameter dynamics estimated from the line scans. *P_c_* and *P_n_* reflect the peak prominence and the noise pedestal at the cardiac frequency.

#### 5.3.2 Imaging system and data acquisition

A custom LSCI system was designed to ensure high repeatability, precision and accuracy of longitudinal imaging experiments. Specifically, we aimed to improve the key factors that potentially increase measurement variation in conventional LSCI: inhomogeneous illumination, low coherence degree, varying polarisation and speckle size [54, 55]. To achieve it, we developed a co-axial illumination LSCI system based on a polarising beamsplitter cube. We used a highly coherent, volume holographic grating stabilised laser diode (785 nm, Thorlabs FPV785P) coupled to a polarisation-maintaining fibre to provide stable and highly homogeneous light output at fixed polarisation orientation. The laser was driven by a combined laser diode and temperature controller (CLD1015, Thorlabs). The light was collimated (F810APC-780, Thorlabs) and delivered onto a polarising beam-splitting cube (CCM1-PBS252/M, Thorlabs) via a custom-built adjustable 1:1 Galilean telescope and a stirring mirror. The laser polarisation was adjusted so the beam-splitting cube would reflect nearly 100% of light, directing it to the infinity-corrected objective (Leica 2.5 N Plan, NA=0.7). The stirring mirror and the telescope were adjusted to achieve homogeneous illumination at the objective’s working distance. The light scattered by the object would be collected by the objective and directed to the beam-splitting cube, where cross-polarised light is transmitted onto the infinity corrected tube lens (TTL200-B) and then to the CMOS camera (Basler aca2040-90um NIR, 5.5×5.5 *µm*^2^ pixels). A neutral-density filter was inserted in the illumination arm to ensure the average light intensity on the sensor would be *≈* 35% of the saturation when recorded at 5000*µs* exposure time [54]. After the initial adjustments, every system component was locked, ensuring no deviations throughout the study. We also performed calibration checks by measuring the coherence degree with a static phantom several times during the project to ensure no changes occurred in the system. In each experiment, LSCI recordings lasted for 600 seconds and were taken at a frame rate of 194 frames per second and an exposure time of 5000 *µs*. Due to the data storage capacity, we limited the field of view to limited to 1024×512 pixels, *≈* 0.5 x 1 mm, therefore producing *≈* 58.2*GB* of data per recording and *≈* 10.5*TB* in total.

#### 5.3.3 Data analysis

LSCI pulsatility analysis is comprised of two essential steps. First, the representative cardiac cycle has to be estimated. Second, vessel-type specific ROIs have to be placed, and vessels have to be dynamically segmented to extract the pulsatility features. We estimate the representative cardiac cycle with the following steps:

1. Mask the field of view to exclude artefacts and areas outside of the cranial window
2. Calculate the average spatial contrast for each frame according to the mask to generate a single refined time course of contrast across the entire field of view. Spatial contrast analysis was described in detail before [56–58]. Briefly, the contrast for each pixel was calculated as 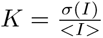, where *σ*(*I*) and *< I >* are the standard deviation and mean of intensity in the 5×5 neighbourhood surrounding the pixel. The contrast calculated for individual pixels, except those masked out in the first step, was then averaged across the frame to produce the average contrast time course. In the present study, each time course contained 116400 elements, corresponding to 194 frames per second for 600 seconds.
3. Identify individual cardiac cycles in the average contrast time course. More specifically, we first applied the Fast Fourier Transform to identify the dominant frequency of the cardiac cycle, with which we determined local minimums corresponding to the individual cycles while allowing some degree of heart rate variability.
4. Characterise each cycle according to a set of features, including but not limited to average, standard deviation, magnitude, duration, the difference between contrast values at the beginning and the end, and the number of noise spikes (increases during the descent phase or decreases during the ascending phase).
5. Reject cycles that do not fit the data quality criteria for the abovementioned features. For most features, the rejection criterion was defined as a deviation from the median value by more than two standard deviations, providing a robust quality control measure.
6. Interpolate contrast images calculated for each cycle, such as every cycle will contain the same number of frames defined as median cycle duration multiplied by the chosen interpolation factor. It is used to avoid feature loss caused by variations in the cycle duration, effectively resulting in irregular sampling and allowing a “super-resolution” temporal analysis approach. It is important to note that only a minor duration variation is permitted in non-excluded cycles, which should not arise from significant changes in heart activity.
7. Average interpolated cycles to produce representitative contrast cycle *K_cycle_* and then convert it to blood flow index as *BFI* = 1*./*(*K_cycle_.*^2^)

From the resulting BFI images, we calculated the per-pixel pulsatility index as *PI* = (*BFI_max_ − BFI_min_*)*/ < BFI >*. Examples of average BFI image and corresponding PI image are shown in Fig. 7, B, C, and Supplementary Figures A1, A2. Such images were then used to guide semi-automated segmentation, where the user selects regions of interest (ROIs) containing segments of arteries, veins or parenchyma. Arterial and vein ROIs are then used as an input for the dynamic segmentation algorithm, modified from our previous studies [49, 51], providing diameter and intra-vessel *BFI* estimations for each frame. Briefly, the algorithm uses an average *BFI* image of the provided ROI to determine the approximate location of the vessel’s centerline and its orientation. Based on it, the average *BFI* profile is calculated across the centre line. Similar to the calculation of the representative cardiac cycle above, the centre line is not perfectly aligned with pixels, effectively resulting in irregular sampling. Therefore, using interpolation during the average profile calculation allows a spatial “super-resolution” approach and retention of the “sub-pixel” information [49]. The values in the average profile are then classified into vessel or background using the minimum intra-class variation approach. Diameter and intra-vessel *BFI* at every frame are then calculated accordingly (Fig. 7, E, G). Parenchymal ROIs are defined as 5000 pixels-sized regions between resolvable vessels and, therefore, do not require dynamic segmentation. Instead, the respective pixel values were directly used to calculate the ROI-averaged BFI (Fig. 7, F).

Finally, following the described above pre-processing, the respective ROI- and vessel-averaged BFI values, as well as vessel diameter values were used to calculate corresponding BFI pulsatility index *PI_BF I_* = (*BFI_max_− BFI_min_*)*/ < BFI >* and diameter pulsatility index *PI_D_* = (*d_max_ − d_min_*)*/ < d >*. These four parameters (*< BFI >*, *< D >*, *PI_BF I_* and *PI_D_*) were used to provide an in-depth characterisation of microvascular pulsatility in the ageing mouse cortex presented in the Results section.

### 5.4 Two-Photon Microscopy

Following the last LSCI recording, we used two-photon microscopy (TPM) to assess capillary density, diameters and pulsatility in 9 out of 12 mice (3 in each age group). All TPM recordings were performed in awake mice. A tail vein catheter for infusion of fluorophores was placed under anaesthesia before the imaging session. Anaesthesia was induced with 3% isoflurane in 0.8L/min oxygen and maintained at 1.5-2% during catheter insertion.During anaesthesia waning, mice were fixed to the imaging stage to prepare for awake-restrained imaging. Imaging was performed on an Investigator-IV two-photon system (Bruker Corporation, Billerica, MA, United States) with PrairieView software version 5.5 (Bruker Corporation). A load of 300*µl* of 0.5% (0.5mg/ml) solution of Texas Red 70kDa was administered via a tail vein catheter for vessel visualisation and z-stack acquisition. We acquired the data following two scanning protocols - entire field-of-view scanning to acquire angiograms and cross-sectional line scans over random capillary branches. For angiograms, a 25x objective (Olympus, WD=8mm) was used to acquire z-stacks of *≈*190*µm* (FOV=515×512, 461.8*µm*^2^, laser excitation 950nm, 300 pockels power, 587 GaAsP). For each line scan, we captured 3 to 4 cross-sectional intensity profiles of the same capillary at a rate of 300,000 pixels per second. Line scans were recorded for 30 seconds, with 1-2ms scan line period, depending on the exact number of pixels per line.

TPM-acquired angiograms were reconstructed in ImageJ using bleaching correction, contrast enhancement with normalisation (saturated pixels=0.4%), background subtraction (rolling ball=50), and 3D filtering (median 3D: x=2, y=2, z=2) and presented as Z projected maximum intensity projections (MIP) and 3D renderings (3D viewer) as shown in Fig. 7, H. Capillary vessels density was then calculated using Deepvess. Firstly, contrast is enhanced in each image in the z-stack and corrected for any motion present. Then, the 3D skeleton of the vessels is extracted and the volume of each vessel segment and the entire angiogram is calculated. Cros-sectional line scans were used to estimate capillary diameters and pulsatility. Due to the specifics of the TPM data, capillary pulsatility analysis has been performed differently from LSCI and included the following steps:

1. Get a rough estimate of capillary boundaries, average intensity and signal-to-noise ratio over the entire scan.
2. Apply the estimates obtained from the previous step to classify pixels as foreground (capillary) and background at every time point using k-means segmentation. An example of the segmented vessel is shown in Fig. 7, I.
3. Calculate coordinates of vessel boundaries from the mask and estimate the capillary diameter as a difference between boundary coordinates. Exclude time-points sections where the estimated coordinates have abnormally deviated from the mean due to either segmentation failing, motion artefact or low signal due to passing red blood cells. The average capillary diameter is calculated at this step by averaging non-excluded diameter values.
4. Use the Lomb-Scargle periodogram [59] to calculate the power spectrum of fluctuations in diameter and boundary coordinates of the vessel. The periodogram is calculated with a 5-second window for frequencies from 1 to 30 Hz and then averaged across the entire observation period and all cross-sections belonging to the same capillary. An example of the resulting power spectrum is shown in Fig. 7, J. We use the Lomb-Scargle periodogram as it allows calculating power spectrums of signals with missing data, such as the coordinates and diameter estimates, following the exclusion of artefacts.
5. Characterise pulsatility in diameter and vessel boundaries as signal-to-noise ratio (SNR) at the cardiac frequency. Specifically, we define 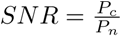 (Fig. 7, J), where *P_c_* is the prominence of the most prominent peak at the frequency range from 9 to 13Hz and *P_n_* is noise pedestal at the same frequency. A noise pedestal is defined as a difference between the peak’s height and prominence; therefore, *SNR >* 1 means that the peak’s power is at least two times higher than the noise.

### 5.5 Isometric wire-myography and histology

Mice were euthanized by cervical dislocation, and the brain was immediately dissected into ice-cold salt solution (PSS in mM: 116 NaCl, 2.82 KCl, 1.18 KH2PO4, 25.0 NaHCO3, 0.03 EDTA and 5.5 glucose; pH 7.4 adjusted at 37*^◦^*C with NaOH and aerated with 5% CO2 in air). Middle cerebral arteries from both hemispheres were gently dissected and mounted in an isometric wire myograph (Danish Myo Technology A/S, Denmark). Following a minimum of 30 min equilibration at 37*^◦^*C in PSS aerated with 5% CO2 in the air, the arteries were normalized in LabChart (ADInstruments, Dunedin, New Zealand). The normalization was based on the construction of passive tension – internal circumference relation, and the internal circumference that corresponds to a pressure of 100 mmHg (IC100) was determined using LaPlace’s equation. 1 The passive wall tension was set to the internal circumference that is 90% of the IC100, i.e., IC1 = 0.9·IC100. It has been shown that resistance arteries develop the maximal contraction at IC1. After 20 min equilibration, the contraction was stimulated three times with thromboxane A2 receptor agonist, U46619 (10-6 M), separated by washouts. Then, the concentration-response curves for U49919 (from 10*^−^*^8^ to 10*^−^*^6^ M) were performed. The myograph experiment data were analyzed in Labchart, following export to Prizm 10 (GraphPad) for further data analyses and statistics.

At the end of the experiment, the arterial segments were fixed directly in the myograph chamber with ice-cold PFA, followed by 24 hours of incubation, and then stored in sterile PBS at 4*^◦^*C until paraphing embedded for histology.

### 5.6 Statistical analysis

Data are shown as means ± SD.

## Data availability

Data underlying the results presented in this paper are not publicly available at this time but may be obtained from the authors upon reasonable request.

## Appendix A Extended data

### Supplementary Figure 1 - average BFI images

**Fig. A1.**
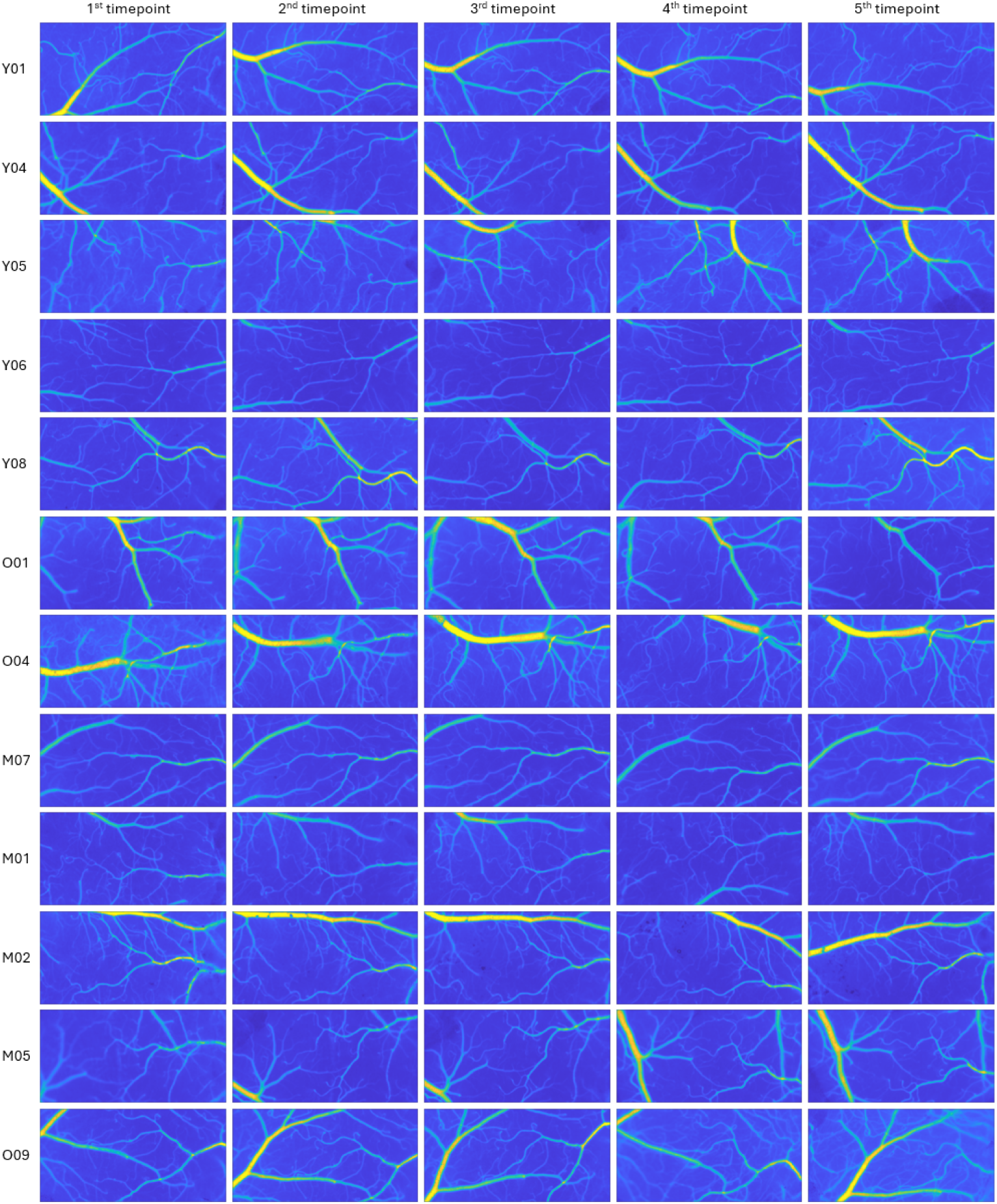
Average blood flow index images for all awake recordings - columns correspond to different time points, rows to different animals. The colourmap limits are identical in all images.

### Supplementary Figure 2 - average PI images

**Fig. A2.**
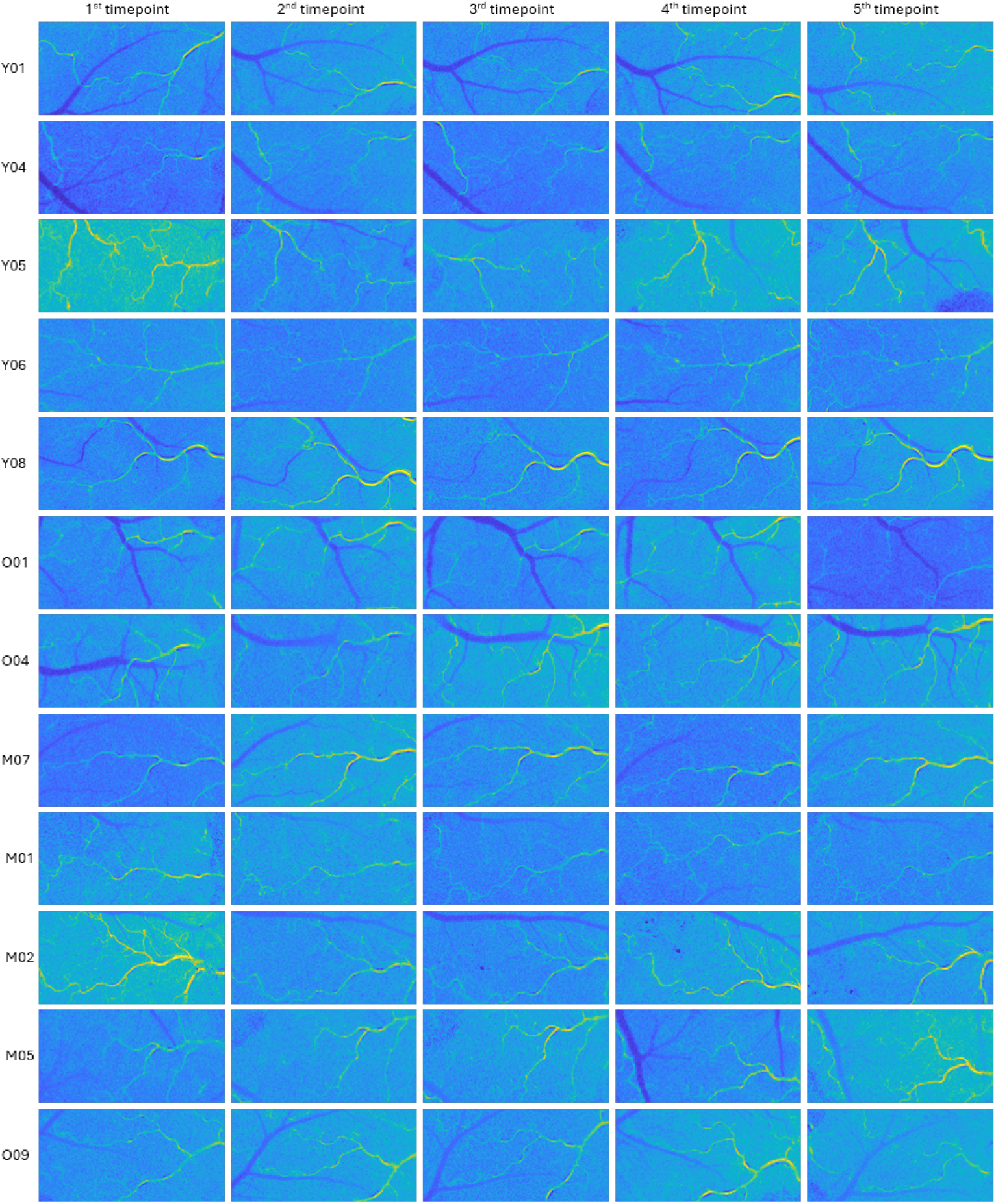
Average pulsatility index images for all awake recordings - columns correspond to different time points, rows to different animals. The colourmap limits are identical in all images.

### Supplementary Figure 3 - number of animals and regions of interest for each time-point

**Fig. A3.**
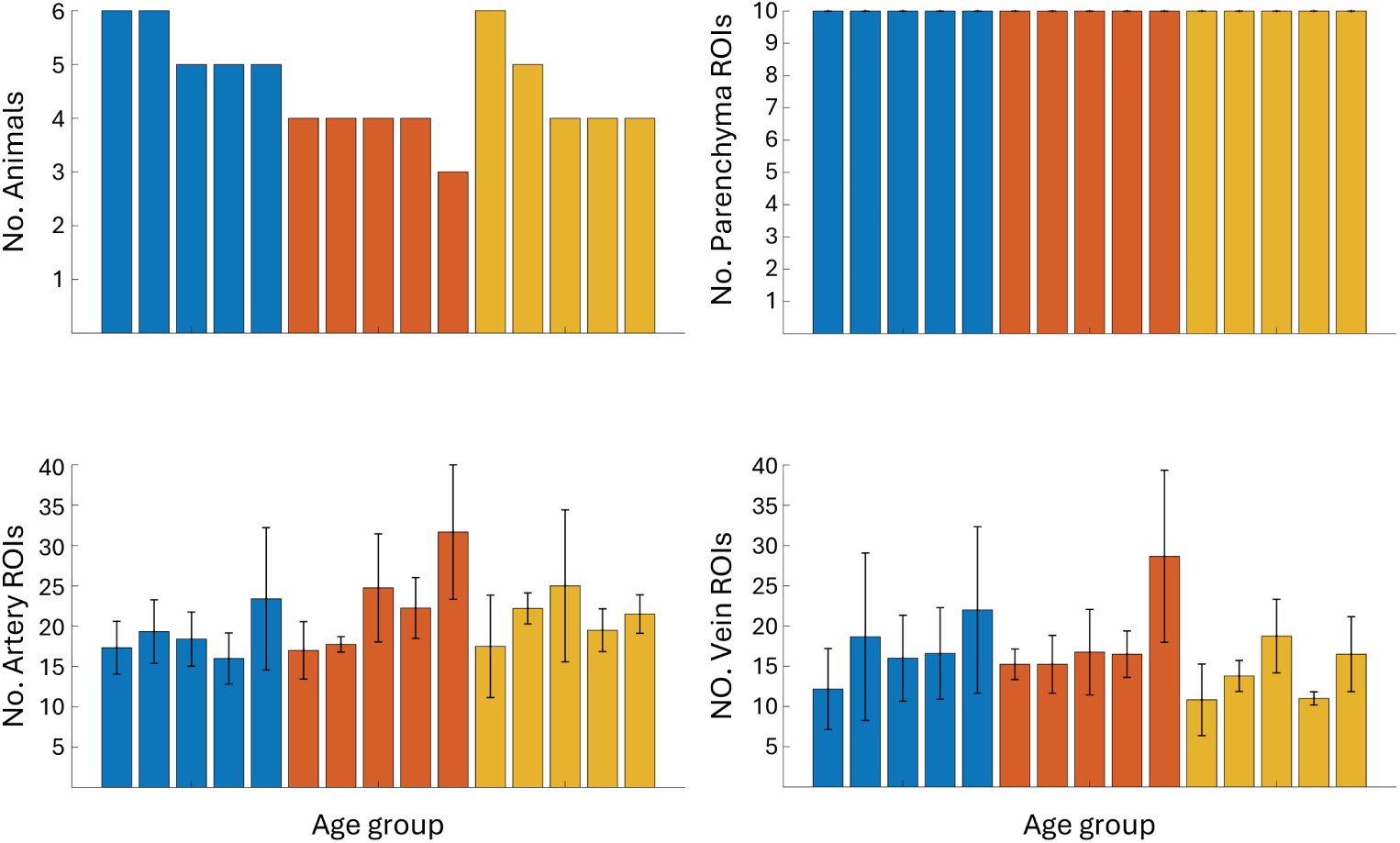
Number of animals and respective regions of interest for each time-point. A – number of animals. Note that the same animals are used for different time-points within the age group. B – number of parenchymal regions. C – number of segmented arteries and arterioles. D – number of segmented veins and venules. E – histogram representing the number of segmented arteries and arterioles according to the diameter across all measurements. F - histogram representing the number of segmented veins and venules according to the diameter across all measurements.

### Supplementary Figure 4 - characteristics correlation in awake animals

**Fig. A4.**
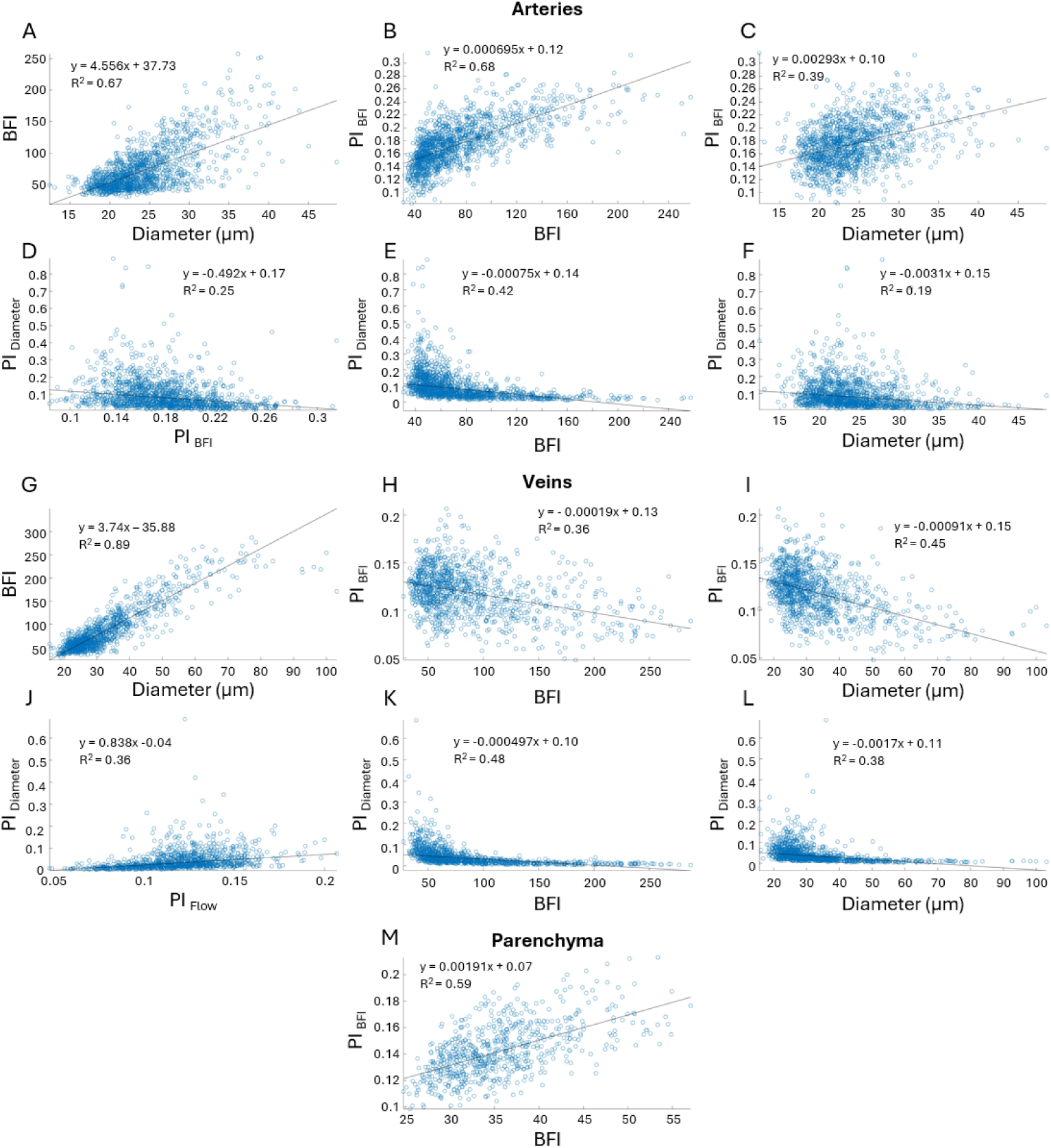
Characteristics correlation based on all measurements in awake animals for different regions of interest. Parameters (*a* and *b*) of respective linear regression models (*y* = *a* + *bx*) and the goodness of fit metric (*R*^2^) are stated in the legend for each graph.

### Supplementary Figure 5 - Periodogram and dynamics examples

**Fig. A5.**
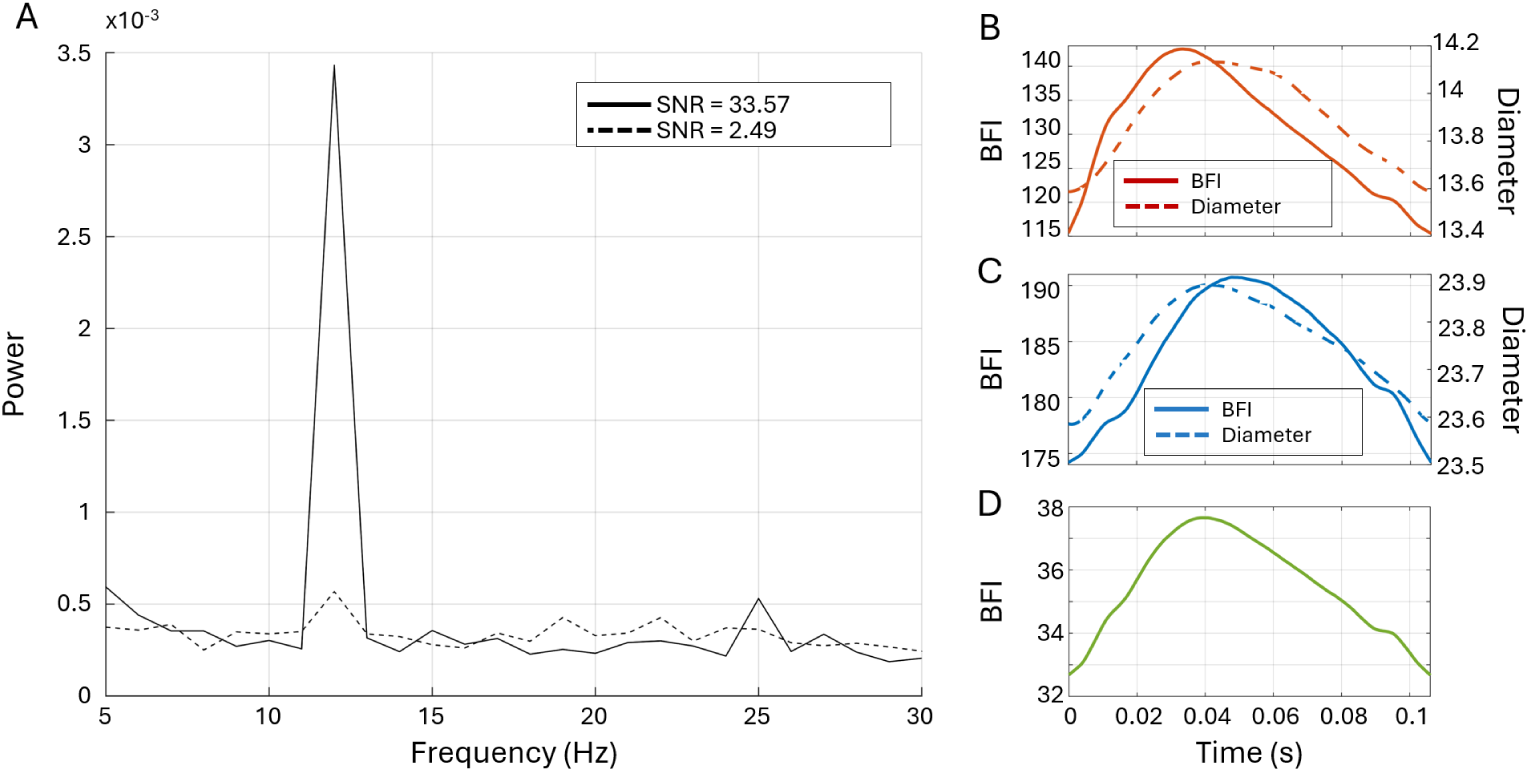
A - examples of Lomb-Scargle periodograms calculated from TPM line scans of strongly and weakly pulsating capillaries. B-D Examples of arterial, venous and parenchymal BFI and diameter dynamics during cardiac cycle.

### Supplementary Figure 6 - Change in pulsatility and perfusion across the microvascular network under anaeshtesia

**Fig. A6.**
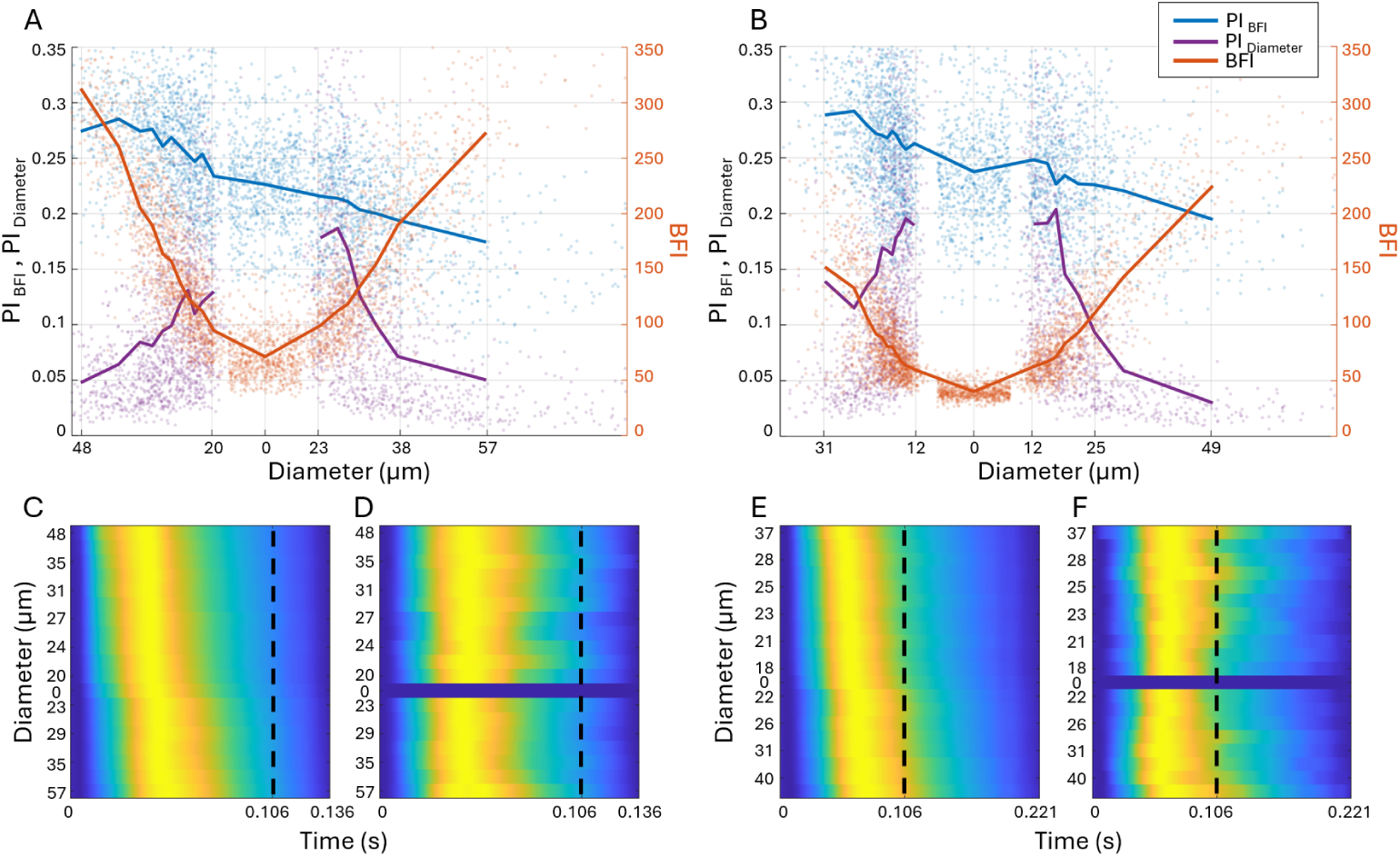
Average diameter and perfusion pulsatilities *PI_D_* and *PI_BF I_* (yellow and blue lines, left Y axis), and blood flow index *BFI* (red, right Y-axis) plotted as a function of the average diameter *D* across the imaged microvascular network. A – during isoflurane anaesthesia. B – during ketamine-xylazine anaesthesia. C, D - respective shapes of the BFI and diameter change during the cardiac cycle under the isoflurane anaesthesia, normalised for each average diameter and presented as a colour-coded map with time and diameter axis. E, F - respective shapes of the BFI and diameter change during the cardiac cycle under the ketamine-xylazine anaesthesia, normalised for each average diameter and presented as a colour-coded map with time and diameter axis. Note how values and shapes have changed compared to the measurements in the same regions of interest in awake animals (Fig. 3). Most interestingly, the decline of the perfusion pulsatility has slowed down – for isoflurane and ketamine-xylazine *δPI_BF I_* between largest arteries and veins is *≈* 0.1, while for awake animals it was *≈* 0.15.

